# HKG: An open genetic variant database of 205 Hong Kong Cantonese exomes

**DOI:** 10.1101/2021.06.15.448515

**Authors:** Min Ou, Henry Chi-Ming Leung, Amy Wing-Sze Leung, Ho-Ming Luk, Bin Yan, Chi-Man Liu, Tony Ming-For Tong, Myth Tsz-Shun Mok, Wallace Ming-Yuen Ko, Wai-Chun Law, Tak-Wah Lam, Ivan Fai-Man Lo, Ruibang Luo

## Abstract

HKG is the first fully accessible variant database for Hong Kong Cantonese, constructed from 205 novel whole-exome sequencing data. There has long been a research gap in the understanding of the genetic architecture of southern Chinese subgroups, including Hong Kong Cantonese. HKG detected 196,325 high-quality variants with 5.93% being novel, and 25,472 variants were found to be unique in HKG compared to other Chinese populations (CHN). PCA illustrates the uniqueness of HKG in CHN, and IBD analysis revealed that it is related mostly to southern Chinese with a similar effective population size. An admixture study estimated the ancestral composition of HKG and CHN, with a gradient change from north to south, consistent with their geological distribution. ClinVar, CIViC and PharmGKB annotated 599 clinically significant variants and 360 putative loss-of-function variants, substantiating our understanding of population characteristics for future medical development. Among the novel variants, 96.57% were singleton and 6.85% were of high impact. With a good representation of Hong Kong Cantonese, we demonstrated better variant imputation using reference with the addition of HKG data, thus successfully filling the data gap in southern Chinese to facilitate the regional and global development of population genetics.

## Introduction

Hong Kong is a densely populated city in southern China. Its population dynamics are strongly associated with its historical background, especially regarding the transition from a British colony into a Special Administrative Region (SAR) of China in 1997, so it has a unique migration history (1). About 90% of the early settlers in Hong Kong are thought to have originated from Guangdong Province in southern China (2), and the majority of them were Cantonese (3). Cantonese is, however, often loosely defined and refers mostly to a Yue-speaking Han Chinese sub-group in the large southern China region. The Yue people also comprise other subgroups, including Teochew and Hakka, possibly with different ancestries. However, all of them are genetically under-represented in current studies and are vaguely regarded as a mixture of southern Han Chinese (4-6). Possibly because of the high population complexity, genetic correlations between subgroups can be found only in a few old studies (6,7). The limited availability of sequencing data in the southern Han Chinese subgroups, including Hong Kong Cantonese, also restricts further investigation into their genetic specificity and connections with other Chinese subpopulations. Therefore, the genetic architecture and composition of the Hong Kong population is still unclear, so there is a need to fill the gap of genetic diversity in Chinese populations.

Several large-scale population genetic studies are available as good references for East Asian and Chinese populations. The Genome Aggregation Database (gnomAD, https://gnomad.broadinstitute.org/) complements the world’s largest variant database dbSNP (8), with variants from Whole-Exome Sequencing (WES) and Whole-Genome Sequencing (WGS) samples from East Asia, providing annotations at the super-population level. The NARD database sampled 1,690 WGS data of several Northeast Asian populations, including samples from Korea, Japan and China (9). Note that only about 3.4% of its samples were obtained from Hong Kong in this study, but not specifically Cantonese. Wang *et al*. compared seven genome-wide data sets from 46 recent groups and 166 ancient groups of East Asians, revealing the evolution of genomic composition in East Asia (10). The worldwide 1000 Genome Project (1KGP) sampled 388 individuals from three well-represented Chinese populations, including Han Chinese in Beijing, southern Han Chinese, and Chinese Dai in Xishuangbanna (7). The GenomeAsia project analyzed WGS data of 1,739 individuals from 219 populations and groups across Eastern and Southern Asia, including China. Recent large-scale Chinese population studies, including ChinaMap and Nyuwa, also involved intensive Chinese sampling (11,12). However, as a representative of southern China, there are very few large-scale genomic resources specific to Hong Kong for characterizing the Cantonese population, which is essential to elucidate its adaptive changes (13) and promote further medical development (14). Owing to the increasing need to gather genetic data for population-wide studies, Yu *et al*. recently analyzed the WES data of 1,116 Hong Kong samples and identified a set of variants for potential pharmacogenetic use in Hong Kong (15). Most of their variant data, however, are not publicly available. The government announced plans to organize a Hong Kong-specific genome institute (HKGP), but it is still at an early stage of development (15). Therefore, our Hong Kong (HKG) database was developed to broaden the availability of freely accessible genomic resources for Hong Kong Cantonese to facilitate more effective intra- and inter-population-wide comparative analyses.

In this study, we describe HKG, the first and by far the largest openly available variant database for Hong Kong Cantonese, extracted using 205 high-quality whole exome sequencing data. Exome sequencing can capture the most informative sections of the entire genome as efficiently as WGS (16), allowing highly confident annotations and effective consequence interpretations. We also show the ability of HKG to better position Hong Kong Cantonese among other Chinese populations, update the population-specific information, especially for clinically significant variants, and improve variant imputation and correlation with local samples. HKG can potentially be a pioneer in providing a reference for development in upcoming regional and local genetics studies.

## Materials and Methods

### Data acquisition

The paired-end 150bp raw reads used in the current study are from the Clinical Genetic Service of the Department of Health of Hong Kong. The samples originated from 205 individuals in the Hong Kong SAR who self-reported as Cantonese. The samples were target captured using SureSelect Human All Exon V6 (Target Size 60M) of Agilent Technologies and were sequenced using Illumina NovaSeq 6000 in the Novogene Tianjin Sequencing Center & Clinical Lab for 200X target depth according to the manufacturer’s instructions.

### Joint variant calling, recalibration and filtering

Cleaned reads were aligned with BALSA (17) to the GRCh38 reference release 5 (GCA_000001405.20) with decoy hs38d1 (GCA_000786075.2), and then sorted with samtools v1.10 (18). Duplicated reads were marked by Picard v2.0.1 (19). Variants were first called from individual samples using the HaplotypeCaller module in GATK v4.1.3.0 (20) and were stored in GVCF format. GenomicsDBImport and GenotypeGVCFs modules were then used to perform joint variant calling on all 205 samples.

The GATK Variant Quality Score Recalibration (VQSR) was used to remove low-quality variants. The SNP VQSR model was trained using HapMap (21) v3.3 and 1KGP Omni v2.5 SNP sites, and the INDEL VQSR model was trained using the Mills et al. 2011 (22) 1KGP gold standard and Axiom Exome Plus indel sites. Sensitivity thresholds of 99.6% and 95.0% were used to filter SNPs and INDELs, respectively. In addition, we filtered out sites using the same filter criteria used by ExAC (23): (1) InbreedingCoeff (inbreeding coefficient) < -0.2; (2) AC (allele count) = 0; (3) DP (depth) < 10; or (4) GQ (Genotype Quality) < 20. The filtration was performed using bcftools (v1.10.2).

### Variant annotation

The stats module in bcftools was used to get the variant statistics. We used bcftools to split multi-allelic records into multiple single-allelic records for variant annotation and conducted annotations of each allele with the Variant Effect Predictor (VEP) tool (Ensembl GRCh38 release 100). dbNSFP4.0a was used to get the annotations of dbSNP ID, GnomAD population allele frequency, ExAC population allele frequency, SIFT, Polyphen2, MetaSVM, MetaLR, CADD, GERP++, phyloP100way, phastCons100way, 1KGP population allele frequency, Exome Variant Server population allele frequency, and ClinVar Clinical significance. ClinVar (v2020-07-06) was used as a custom annotation in VEP to retrieve the pathogenic variants. We used GATK LiftoverVcf to liftover the CIViC (v2020-08-07) database from GRCh37 to GRCh38 before using it to get druggable variants. The LOFTEE (24) plugin of VEP was used to generate the loss-of-function variant annotations. Scripts from (24) were used to retrieve the multi-nucleotide variants (MNVs). We used DAVID (http://david.abcc.ncifcrf.gov) to perform the enrichment analysis of gene ontology (GO) biological processes and KEGG pathways.

### Interpopulation comparison

1KGP Phase 3 (1KGPp3 20170504) variants were downloaded from ftp://ftp.1000genomes.ebi.ac.uk/vol1/ftp/release/20130502/supporting/GRCh38_positions/. A Chinese-only dataset was generated by extracting the samples labeled CHB, CHS, or CDX (referred to as 1KGP CHN in the following) and excluded variants with AC = 0. These variants were then merged with HKG by bcftools (v1.10.2). Variants at positions either without the ‘GRCH37_38_REF_STRING_MATCH’ tag (25) or not in the SureSelect Human All Exon V6 bed regions were filtered out. The R package gdsfmt was used to convert the datasets into gds format for SNPRelate to use as input for PCA analysis (26,27).

For the IBD analysis, variants in HKG were phased using Beagle 4.1 (28,29). The 1KGP phase 3 variants (1KGPp3 20170504) were used as the reference panel. All HKG and 1KGP CHN variants were processed by the Beagle v4.1 IBD calling algorithm with 15 iterations, each time using a different random seed. In each iteration the minimum length of the IBD was set to 3 cM to minimize the influence of phasing and genotyping errors. We combined the results of the 15 iterations using the “ibdmerge” module. The normalization method from Nakatsuka et al. 2017 (30) was used to enable a population-wise comparison.

For admixture analysis, we aggregated the variants in the HKG exome bed regions, 1KGP CHN, other 1KGP EAS (i.e., JPT, CHB and KHV), and SAS (GIH, PJL, BEB, STU, and ITU). We used PLINK (v1.90b6.10 64-bit) (31) to remove the variants with (1) missing call frequencies greater than 0.05, (2) with minor allele frequency lower than 0.05, or (3) with Hardy-Weinberg equilibrium exact test p-values below 0.0001 (--geno 0.05 --maf 0.05 --hwe 0.0001). For each window of 1,000 SNPs, we calculated the Linkage Disequilibrium (LD) between each pair of SNPs in the window and filtered those LD greater than 0.2. The filtration was repeated by moving 100 SNPs forward until all SNPs were scanned (--indep-pairwise 1000 100 0.2). After the filtrations, ADMIXTURE v1.3.0 (32) was applied to the rest of the variants, with the estimated number of subpopulations (*k*) ranging from 2 to 15. The output of the ADMIXTURE was visualized by Pophelper v2.3.0 (33).

### Enrichment analysis of novel variants

The mapped genes from the high-impact novel variants of HKG were obtained, as well as the three CHN populations of 1KGP, namely CHS, CHB, and CDX. DisGeNet (version 7.0) data was used to identify the gene-disease associations. The significant enrichments of those non-overlapped genes among the four populations in HKG were estimated by R package “phyper”. The overlap between the disease-gene sets of DisGeNet and the 634 HKG uniquely affected genes (i.e., those not contained in the mapped genes from CHN) were used to calculate the p-values of the enrichment. Only enrichments with p < 0.01 and FDR < 0.25 were selected.

### Imputation and correlation analysis

To verify the improvement of the imputation accuracy using the HKG data as a local reference panel in addition to the 1KGP samples, two imputations on autosomes were performed: (1) using only the 1KGP samples as the reference panel (1KGP), and (2) using both 1KGP and HKG samples (1KGP+HKG). We randomly divided the 205 HKG samples into two sets, one with 204 samples used as the reference panel, and the other with one sample used as the test data. This step was repeated five times to generate five reference-test pairs for 5-fold cross validation. The 204 samples of each pair were merged with the 1KGP samples to create the 1KGP+HKG reference panels. Variants with AC < 3 or (AN - AC) < 3 or with missing genotypes were excluded. Only biallelic variants were kept for analysis. For each test sample, we obtained the imputable variant intersection of the 1KGP and 1KGP+HKG. In each chromosome, we randomly masked 200 intersected variants in the test samples. The imputation by BEAGLE 4.1 using the default setting along with niterations=10 and ne=20,000 was performed on each test sample with the two reference panels (1KGP and 1KGP+HKG). To assess the imputation quality, the info score for each imputed variant was calculated. An imputed variant with info score < 0.4 (resp. > 0.7) was classified as “poor” (resp. “confident”). The change in info scores after the addition of HKG samples were further compared under different MAF ranges.

To assess the performance of HKG for correlation analysis using local samples, 58 whole genome sequencing Hong Kong individuals from the Northeast Asian Reference Database (NARD;(9)) and 532 pharmaceutical-related variants from Yu *et al*. 2021 (15) were used. The NARD dataset (NARD_MAF.hg38.vcf.gz) were downloaded from https://nard.macrogen.com/, and 17,224 variants having AC larger than 1 in its Hong Kong samples were considered. The 532 variants from Yu *et al*. 2021 were liftover from GRCh37 to GRCh38 using GATK LiftoverVcf. Variant intersection was obtained using bcftools isec. Variant allele frequency regression and calculation of Cook’s distances were performed using the statsmodels module in Python.

## Results

### Discovery of variants using the WES of 205 Hong Kong Cantonese individuals

Deep whole exome sequencing obtained data from ∼38Mbp targeted positions (92.24% exomic, 4.29% intronic, 3.47% others such as intragenic positions). The average on-target mean depth was ∼159x (96-277x per sample), with 85% of target regions covered by on average of at least 52x (24-101x) and an average of 64% (40-85% per sample) covered by at least 100x. An in-house pipeline was used to process and interpret the WES data (Supplementary Figure S1, also see Materials and Methods). We found 196,325 high-quality variants (83.99% of all called variants), including 186,466 SNPs, 3,709 insertions, and 6,150 deletions, after intensive quality filtering. On average, 26,221.6 variants were called per sampled individual. The transition/transversion ratio (Ti/Tv) was 2.89 for all bi-allelic variants that passed quality control. Breaking down by minor allele frequency (MAF), the Ti/Tv ratio for variants with MAF > 5% and MAF ≤ 5% were 2.93 and 2.87 respectively. All variants were categorized based on their AC in HKG (Figure 1A). Compared with the worldwide distribution available in gnomAD among the HKG targeted region, both showed approximately half of the variants as singletons (AC = 1). We found that HKG had more common variants (i.e., AC > 10), possibly reflecting lower genetic diversity in the population or population-specific variants that exert positive selection (34).

**Figure 1.**
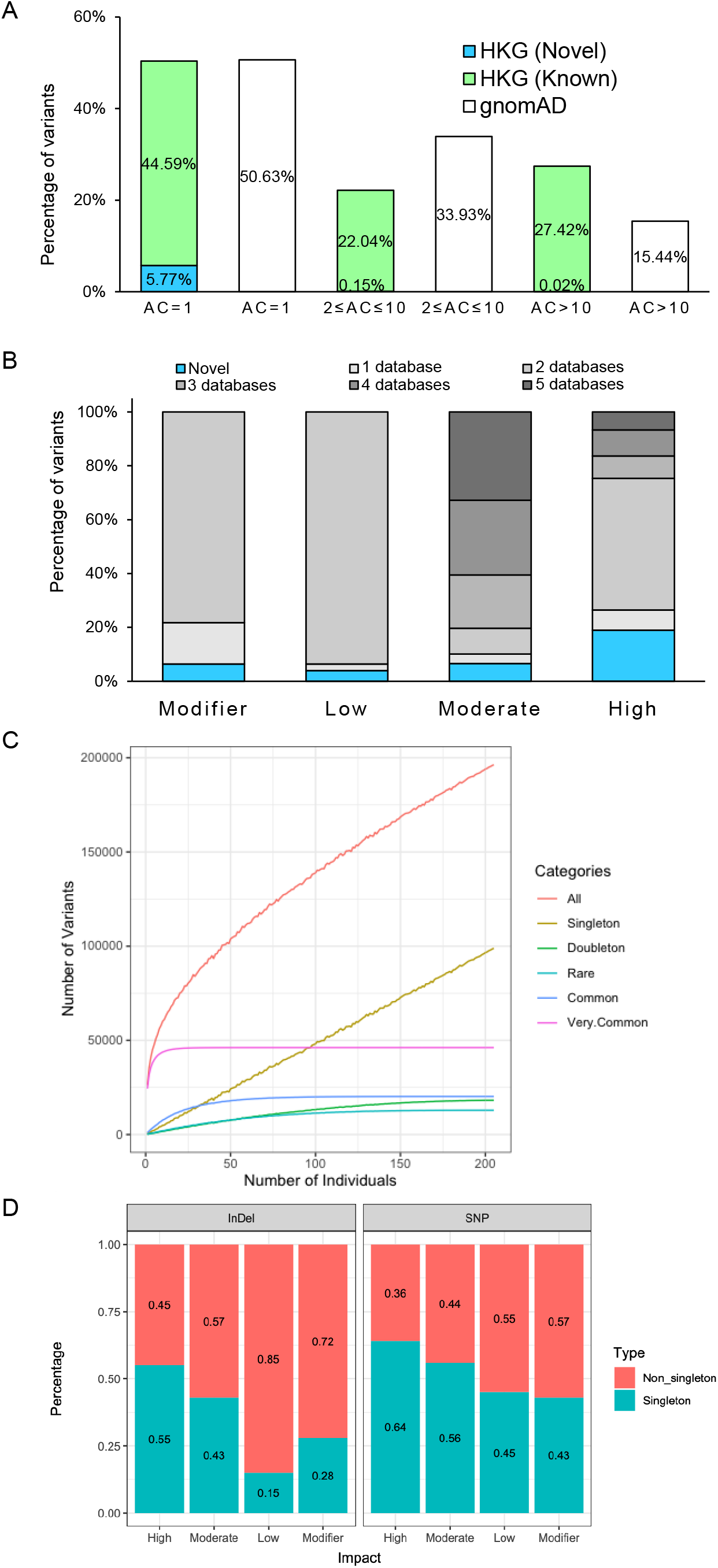
Variant compositions of HKG samples. (A) Comparison of variants in HKG and gnomAD under different allele counts. (B) Number of variants as a function of the number of individuals. (C) Percentage of novel variants and known variants according to MAF categories. (D) Percentage singletons of different variant types and impacts.

Using the Variant Effect Predictor (VEP) of Ensembl, the identified variants were annotated with different consequences. Five main public databases (dbSNP151, 1000 Genomes, ESP6500, ExAC, gnomAD) were used to classify each HKG variant as either known (annotated in at least one database) or otherwise novel. 93.85% of known HKG variants were annotated in at least two databases; 88.54% of HKG singleton variants and 98.87% of HKG doubleton variants were known (Figure 1B). The relative contribution of annotations from each database is: 36.18% dbSNP151, 33.93% gnomAD, 13.72% ExAC, 10.03% 1KGP, and 6.14% ESP6500. There were 11,659 novel HKG variants, with over 96.57% of them being singletons. Since the variants were confidently identified using GATK joint variant calling, the singleton variants were likely true variants rather than errors. The large proportion of novel singleton variants indicates that our analysis using high-depth WES data was highly sensitive. But the number of singletons detected increased with the addition of samples (Figure 1C) suggested that the sampling is not saturated.

High-impact variants are defined as potentially altering the protein structure, and therefore affecting functionality. HKG variants were grouped into four categories by their function impact on a transcript or coding genes in decreasing severity: (1) HIGH, including stop-gain or stop-loss variants, frameshift variants, splice donor or acceptor variants and initiator codon variants, (2) MODERATE, (3) LOW, and (4) MODIFIER (Table 1). The variants (SNPs or INDELs) of higher impact were more likely to be singleton than those of lower impact (Figure 1D). A large number of variants were missense (42.46%), synonymous (31.32%) or intron (8.81%) variants. However, there were relatively few missense mutations at high MAF: at MAF > 1%, the distribution became 15.26% missense, 23.99% synonymous, 23.50% intron. On the other hand, deleterious mutations, such as frameshift, gain of stop codon, and splice site variants, were mostly singletons. INDELs were generally less common as they are likely to cause deleterious frameshift, so many more singletons were found as SNP in the HKG exome data (Figure 1D). There was a lower proportion of INDEL singletons with LOW and MODIFIER impact (15% and 28% resp. in INDEL; 45% and 43% resp. in SNP; details in Table 1) as well.

**Table 1.**
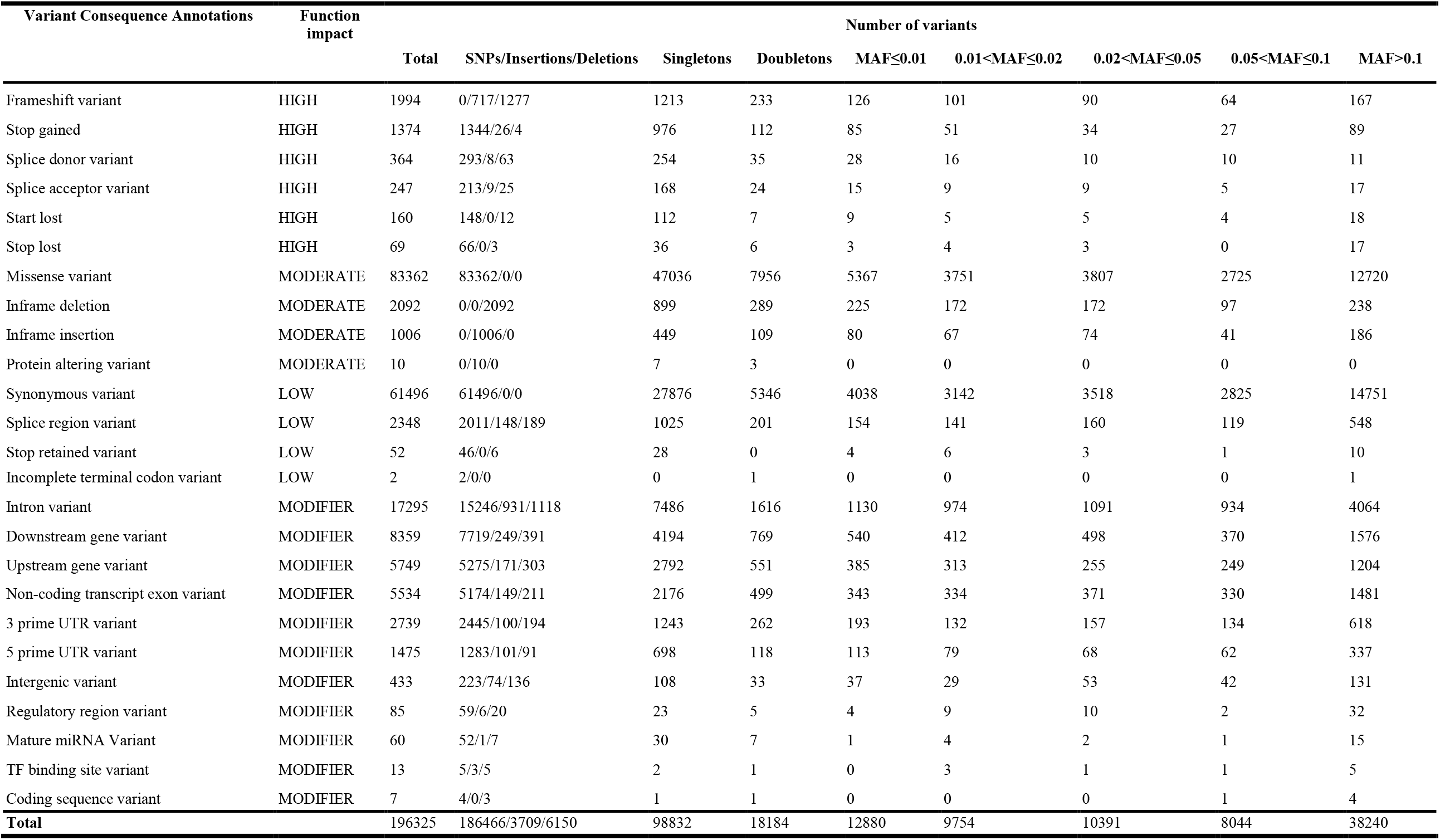
The count of variants in HKG by function impact and MAF.

A multi-nucleotide variant (MNV) is a combination of multiple variants coexisting in the same codon on the same haplotype. There were 800 MNVs that occurred in at least two individuals, and 254 of them were caused by two consecutive nucleotide changes. Since exome regions have higher GC content than other regions (35) and CpG sites have higher mutation rates (30), we observed an expected high percentage (36.81%) of MNVs related to transitions at CpG sites (Supplementary Figure S2). We found 658 (82.25%) MNVs that might alter the protein in a different way than considering two single variants separately (36). The alteration of consequence is listed in Supplementary Table S1. 48.88% of the MNVs were missense variants. All nonsense single variants found in MNV pairs were rescued in HKG, so they were no longer causing protein truncation or disease. Rescued nonsense MNVs were involved in disease-related genes *COL9A2, SYNE2* and *DNAH11* (37). *COL9A2* is associated with autosomal dominant Multiple Epiphyseal Dysplasia 2, and *SYNE2* is related to autosomal dominant Emery-Dreifuss muscular dystrophy 5. *DNAH11* is associated with autosomal recessive primary ciliary dyskinesia-7. Severely affected consequences, such as “gained nonsense” and “rescued nonsense” MNVs, were found in various genes. Among the gained nonsense MNVs, one of the affected genes was *HLA-DRB1*, which is related to rheumatoid arthritis. The variants chr6:32584314G>T and chr6:32584315A>T in *HLA-DRB1* were originally classified as separate missense variants, while their MNVs was identified as a nonsense mutation (i.e., changing the Phenylalanine to a stop codon). This MNV was heterozygous in 4 HKG samples, affecting only one copy of the gene. We found two other heterozygous nonsense MNVs in *HLA-DRB5* (chr6:32522110GA>TT) and *SLC30A8* (chr8:117172544CG>TA), in 4 and 12 individuals respectively. We found a stop-loss MNV chr14:105907227TT>CC in *IGHD2-8* shared by 33 individuals, and none of them are homozygous. *IGHD2-8* is one of the *IG D* (diversity chain immunoglobulin gene) that undergoes somatic recombination before transcription. During somatic recombination (38), a pair of *IG D* and *IG J* genes (joining chain immunoglobulin gene) are joined together by randomly deleting the DNA between them. After somatic recombination, somatic hypermutations in the mRNA can also increase immunoglobulin diversity. The random somatic recombination and somatic hypermutations could dilute the impact of this MNV by deleting or changing the sequence that contains the MNV. The variant chr11:71816849CG>TA on *ZNF705E* was homozygous, found in 12 individuals, which might introduce an early truncation to the protein. It was also located at the last exon of the gene with only one transcript, and therefore might have functional implications.

### Population comparison of HKG variants

The 1KGP project contains the largest amount of publicly available Chinese (CHN) population genetic data, including southern Han Chinese (CHS), Beijing Han Chinese (CHB) and Dai minority in southwestern China (CDX). Other recently developed Chinese databases (11,12) do not provide the full variant list for batch download, so they were not included for HKG comparison. There were 128,470 variants recorded among the four populations. There were 25,472 variants uniquely found in HKG compared with CHN, and 4,366 shared variants with 5-fold differences in at least one CHN. The CHS has the highest number of shared variants with HKG (81,658 shared variants), while CHB has the largest number of unique variants shared with HKG (7,673 variants) (Figure 2A). All four populations had a similar percentage of singletons. Our PCA analysis suggests a unique composition of HKG among CHN, even compared with the closely related CHS, although there was no clear separation boundary among CHB, CHS and HKG (Figure 2B). This suggests that the genetic relatedness of CHN and HKG has been correlated with their geographic location. In addition, as expected, we confirmed that HKG was part of the East Asian (EAS) population by PCA (Supplementary Figure S3).

**Figure 2.**
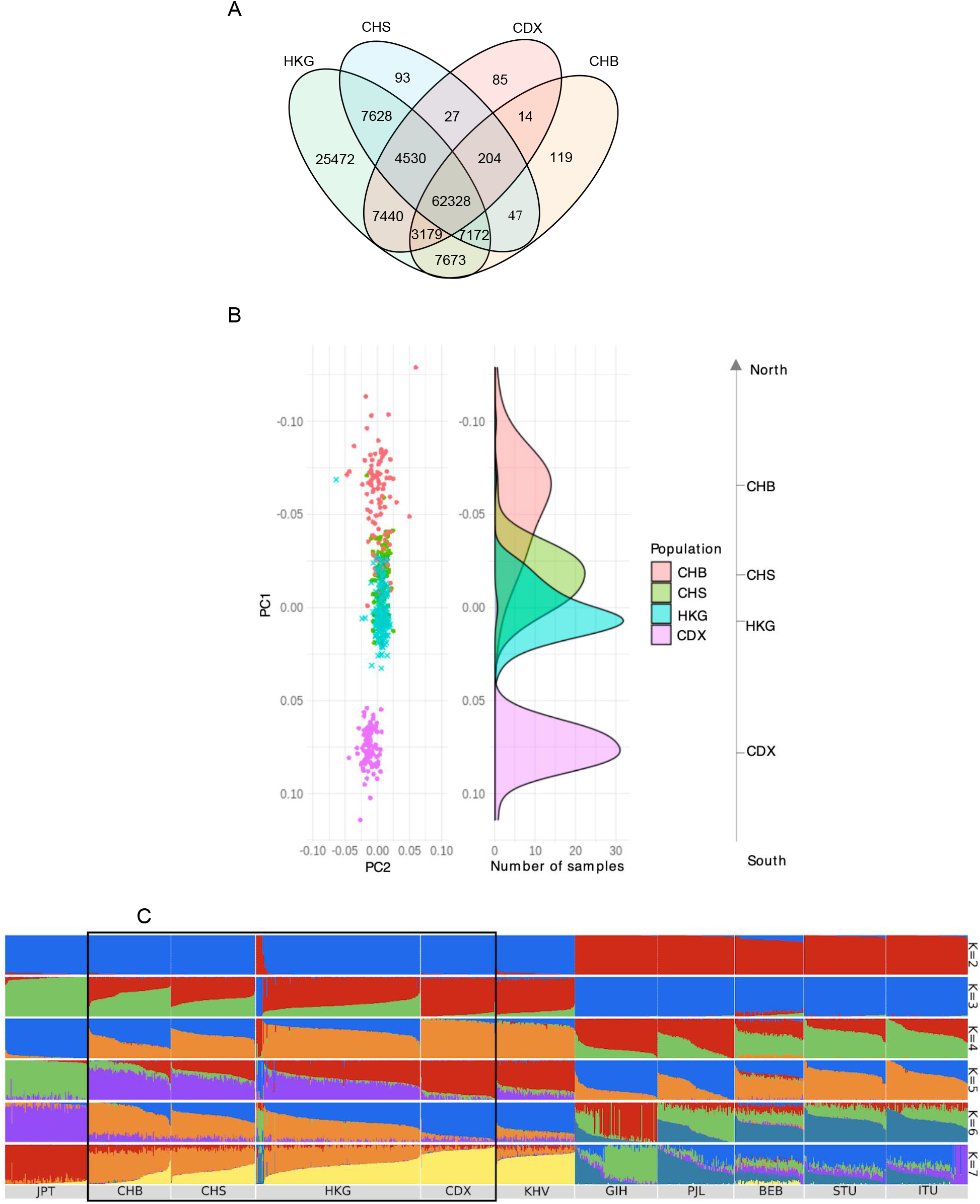
Comparison of variants among HKG and other populations. (A) Venn diagram of variants in HKG and three Chinese populations of 1KGP CHS, CHB and CDX. CHB: Han Chinese in Beijing, China; CHS: southern Han Chinese; CDX: Chinese Dai in Xishuangbanna, China. (B) PCA of HKG and CHS, CHB and CDX. (C) ADMIXTURE analysis of HKG samples with East Asian and South Asian samples in 1KGP (k ranges from 2 to 7). Number of ancestries K = 5 best fits the model. Different colors represent different ancestry components. JPT: Japanese in Tokyo, Japan; KHV: Kinh in Ho Chi Minh City, Vietnam; GIH: Gujarati Indian from Houston, Texas; PJL: Punjabi from Lahore, Pakistan; BEB: Bengali from Bangladesh; STU: Sri Lankan Tamil from the UK; ITU: Indian Telugu from the UK.

Identical-by-Descent (IBD) analysis using the phased variants of CHN and HKG to estimate the effective population size and understand the population demography (39). We analyzed 593 phased samples (CHB: 108; CDX: 109; CHS: 171; HKG: 205) and detected 2,966 IBD segments. For 94 CHB, 93 CDX, 105 CHS, and 205 HKG individual samples with the IBD segment over 3 cM, each individual share at least one IBD segment with 3.51, 6.02, 9.28, and 18.34 individuals on average. These individuals are defined as “relatives” (40). The larger number of relatives indicated a closer relationship among those individuals within the population. Table 2 summarizes the number of detected IBD segments, their total length, and the corresponding number of individuals in each population. CDX had the highest normalized IBD shared and was 10 times higher than that in CHB. A smaller normalized length of IBD in CHB might be due to its large effective population size or diverse sampling. The limited effective population size detected in CDX also agreed with the fact that CDX was a geographically isolated population (in Xishuangbanna of Yunnan province in southwestern China). Having a similar normalized per-segment length, HKG has a level of genetic relatedness like CHS. This suggests that the HKG is not as isolated as the CDX and has a lower population mixture than the CHB.

**Table 2.** Identical-by-Descent (IBD) analysis results of HKG and 1KGP Chinese populations.

| Dataset (Number of samples) | 1KGP_CHB (108) | 1KGP_CHS (171) | 1KGP_CDX (109) | HKG (205) |
| --- | --- | --- | --- | --- |
| Aggregated total length of IBD segments | 899,426,114 | 4,235,340,758 | 9,811,520,175 | 5,537,975,910 |
| Normalized length of IBD segments | 38,916 | 72,847 | 416,731 | 66,212 |
| Number of IBD segments | 167 | 514 | 1299 | 986 |

To investigate the ancestral population structure of HKG, we performed an ADMIXTURE analysis of CHN, HKG, five EAS populations, and six South Asian populations using 1,159,511 autosomal markers. The lowest cross-validation error was achieved when the number of hypothetical ancestral components was set to 5 (K = 5), as shown in Figure 2C with full illustration in Supplementary Figure S4. With two hypothetical ancestral components (K = 2), the results did not show a big difference between HKG and CHN, but HKG also captured a few individuals with completely different major ancestral components than the rest of HKG. These outliers consistently (from K = 2 to K = 15) showed a similar composition to other East Asian populations, such as Punjabi of Pakistan, and Bengali of Bangladesh, which were common minorities in Hong Kong for generations (41). K = 5 showed a clear separation of the populations, with three ancestral components dominated in HKG and CHN (in Vietnam as well). Two of the major ancestral components gradually changed in proportion from the northern to southern Chinese subpopulations. The dominant ancestral populations in the northern Chinese (purple) were progressively diluted towards the south, whereas the proportion of the minor ancestral population in the northern Chinese (red) gradually increased towards the southern Chinese. The compositions of these two major components also showed differences between the closely related CHS and HKG, suggesting deviation in the ancestry of Hong Kong Cantonese from CHS. Further separation from K = 7 onwards becomes ambiguous, suggesting a simple ancestral population in China, which agrees with other Chinese population studies (11,12).

### HKG-specific clinical annotations

A total of 96,795 variants (49.3% of all high-quality HKG variants) identified in at least two HKG individuals were classified as high-confidence variants, 31.26% of which were rare variants (MAF < 1%). To evaluate the potential contribution of HKG variants for biomedical use, we investigated 189 variants with pathogenicity, 410 druggable variants, and 360 LoF variants in greater detail.

Among the 189 ClinVar pathogenic variants found in HKG, only seven were reported in existing studies with Hong Kong samples (9,15). There were 169 annotated pathogenic rare variants (MAF ≤ 1% in HKG and worldwide) located in 141 genes. The details of these annotated pathogenic variants and their annotations are listed in Supplementary Table S2. Among the 20 annotated pathogenic variants which were common (MAF > 1%) in HKG, were reported as pathogenic by other Chinese population studies in ClinVar. The pathogenicity of these variants among Hong Kong Cantonese should therefore be further evaluated. Among the rest, nine pathogenic variants were found to have much higher AF (> 5 times higher) in HKG compared to the worldwide records, and 13 were also found to be common variants in gnomAD of other EAS samples (Supplementary Table S3). Also, 12 common variants were annotated pathogenic with reference only to Western and East Asian populations studies, so they lack support from Chinese data. Thus, it is reasonable to conjecture that these high MAF variants are not pathogenic in HKG.

In total, 410 pharmaceutical-related high-confidence variants were annotated in CIViC and PharmGKB databases, providing a possible guide for drug use. Only four of them were also reported in a recent pharmacogenetics study (15). Among the 24 CIViC variants in HKG, 22 are common variants (MAF > 1%), so they are more likely to be drug effective in the population. We also found that some of these variants had significantly different MAF between the Chinese and non-Chinese populations. These variants might affect drug usage and treatment plans. For example, the variant rs1799782, which is related to Lung Non-small Cell Carcinoma and can increase the response rate of chemotherapy (42), has a much higher MAF in HKG (30.50%) and gnomAD EAS (30.52%) than in the worldwide distribution (9.46%).

There were 401 HKG variants with PharmGKB annotations, six of which were not found in gnomAD but were common in HKG (MAF > 1%). Two of these variants, rs6318 and rs3758581, had an extremely high MAF of over 95%. The G allele of variant rs6318 accounted for 98.5% in HKG, which could be associated with an increased likelihood of drug-related weight gain based on three different human studies (43-45). Another variant, rs3758581, had 95.1% MAF, which is associated with drug response of fluoxetine, fluvoxamine, and voriconazole based on 3 *in vitro* studies (46-48).

Other than database annotations, we found 1,276 LoF variants that occurred in at least two HKG samples using the LOFTEE annotation in VEP, 878 of which were removed due to low confidence. Three variants were marked as false positives because the alternative allele was the ancestral state (35). Common variants (MAF > 0.5) in gnomAD were regarded as errors in the reference (35) and were not analyzed. Therefore, we found 306 highly reliable LoF variants on 301 genes in HKG; 68.5% of the variants were SNPs and 31.9% INDELs; 175 variants were rare (MAF ≤ 1%) and 185 were common (MAF > 1%). However, there was not significantly enriched GO term, KEGG pathway or disease association being found.

### Analyses of novel variants in HKG

By comparing five promising databases – dbSNP, gnomAD, 1KGP, ExAC and ESP6500 – we obtained 11,659 novel variants (10,641 SNPs / 366 insertions / 652 deletions) after stringent filtering. Out of these novel variants, we identified 26 common variants (MAF > 5%), 88 low-frequency variants (MAF = 1–5%), and 54 rare variants (MAF < 1%). More than 96% of the novel variants were singletons (i.e., AC = 1), which was higher than that in ChinaMAP (75.3%) and NyuWa (86.8%), using whole genome sequencing data. The impact distribution of these novel variants was: 18.99% HIGH, 6.56% MODERATE, 3.94% LOW, and 6.39% MODIFIER. The proportion of high-impact variants in the novel sets is significantly higher than the known sets (Figure 3A). A comparison of novel and known variants in different AF categories under each impact level is shown in Supplementary Table S4.

**Figure 3.**
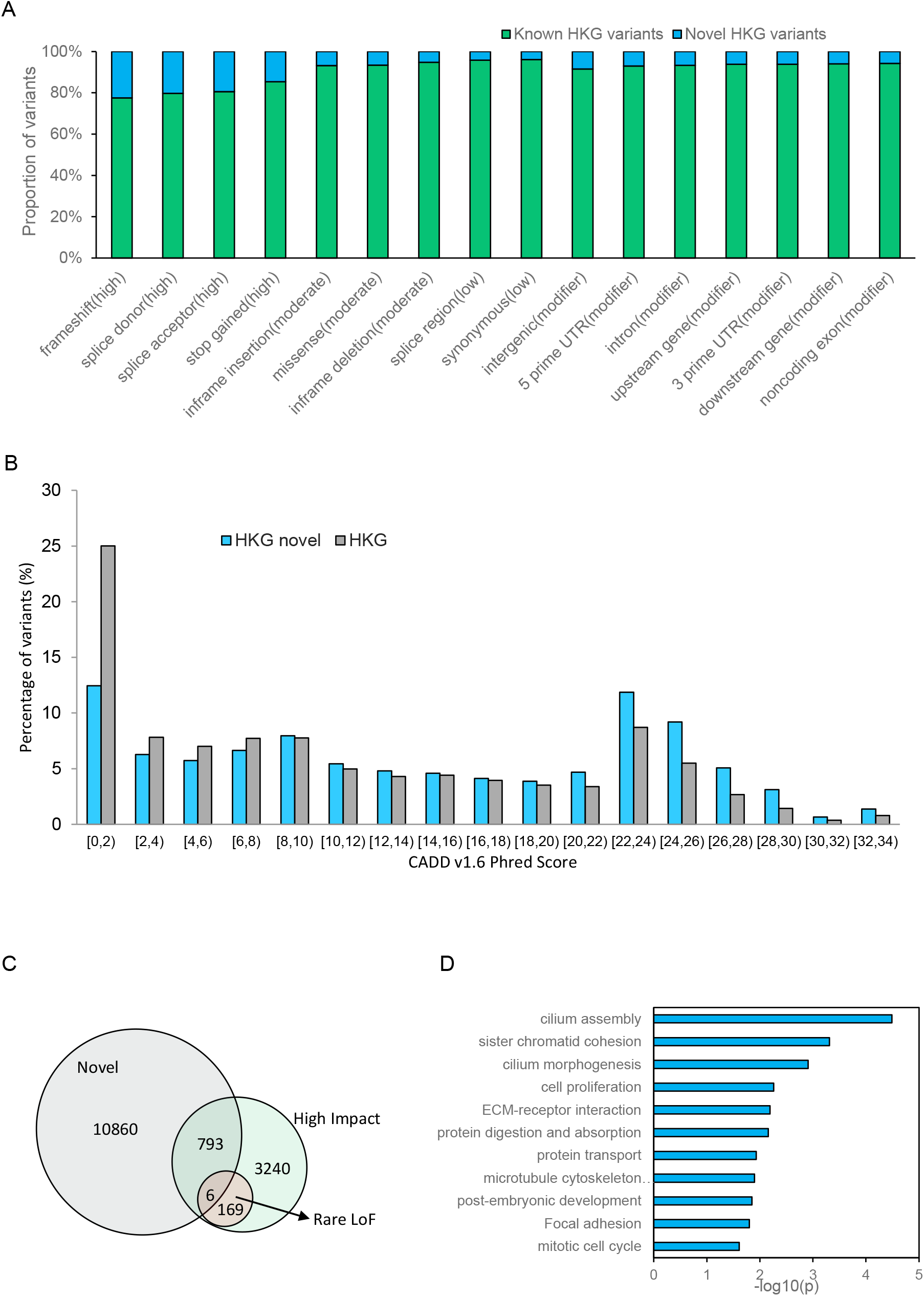
Analyses of the novel HKG variants. (A) Proportion of known and novel variants according to consequences. (B) the pathogenicity score in novel and all variants of HKG. (C) Venn diagram of novel variants, rare LoF and high impact variants. (D) Significantly enriched GO terms and KEGG pathways in 731 genes responsible for the novel high-impact variants of HKG.

We computed pathogenicity scores using the C-scores reported by CADD (Combined Annotation-Dependent Depletion), a widely used database of pathogenicity. A higher C-score or pathogenicity score means more deleterious outcomes. The mean pathogenicity scores of novel variants and all (i.e., novel + known) variants of HKG were 14.4 and 10.5, respectively. A larger proportion of novel variants fell into the high score range (> 20) compared to all variants of HKG (Figure 3B), indicating that a larger percentage of novel variants were more deleterious. Among the 175 rare LoF variants in HKG, six (3.43%) were also present in 799 VEP annotated high-impact novel variants (Figure 3C). The biological association study mapped 799 high-impact novel HKG variants to 731 coding genes, which were significantly enriched with cilium assembly and morphology, ECM-receptor interaction, protein transport, microtubule cytoskeleton organization, sister chromatid cohesion, and protein digestion and absorption (Figure 3D). We also obtained a set of high-impact variant-mapped genes that were present only in HKG but not in other CHN. Other disease-associated genes which were potentially related to high-impact novel variants, based on DisGeNET and their association network, are listed in Supplementary Table S5 and Supplementary Figure S5 for reference. We found eight of the disease-associated genes with novel high-impact variants in at least three individuals (Supplementary Table S6). Our detailed studies suggest that these high-impact novel variants often existed on the same exon of the affected genes. Known pathogenic variants could also be found on the same exons of *MUC4* (49) and *CACNA1A* (50) genes. However, owing to the limited sample size and uncertainty by *in silico* predictions, further confirmation of their pathogenicity is awaited with larger sampling in the Hong Kong Genome Project (HKGP) (51) in the future.

Among the 26 common novel variants (MAF > 0.05), six were in coding regions, although these variants are likely to be benign in HKG given their high AF. The coding region variant chr7:100773854G>GTT of the *ZAN* gene occurs in 7.80% of the population. Deleterious mutations in *ZAN* might affect the adhesion of sperm to eggs, thus reducing fertility (52). There is also a known benign record listed in ClinVar at the same genomic position as the frameshift G>GT variant. Another high AF variant associated with immune response was found on chr7:142796847A>T of the TRBJ2-3 gene, with AF 20.97%. The closely associated pseudogene in the same family chr7:142796707C>G TRBJ2-2P also had a high MAF of 20.20%, but no functional significance was recorded. All the HKG common novel variants are listed in Supplementary Table S7.

### Efficiency of HKG for imputation and correlation

We validated the effective usage of HKG variants using imputation and correlation studies. The imputation accuracy was evaluated using HKG+1KGP and 1KGP reference panels to impute variants in 200 randomly selected positions in five phased Hong Kong Cantonese samples. The addition of HKG in the reference panel yielded a significant increase of 2.189% on the average info score in autosomes compared with using just 1KGP data as panel (Supplementary Table S8). For variants with MAF > 5% and MAF ≤ 5%, the info score increment reached 2.365% and 1.807% respectively. Improvements in imputation quality could be observed as there were 1.33% more high-confidence variants (info score > 0.7) and 1.495% fewer low-confidence variants (info score < 0.4). These improvements were found to be significant, based on the student T-test, marked as ** on Figure 4A, suggesting Hong Kong-specific data should be included in imputation of local samples.

**Figure 4.**
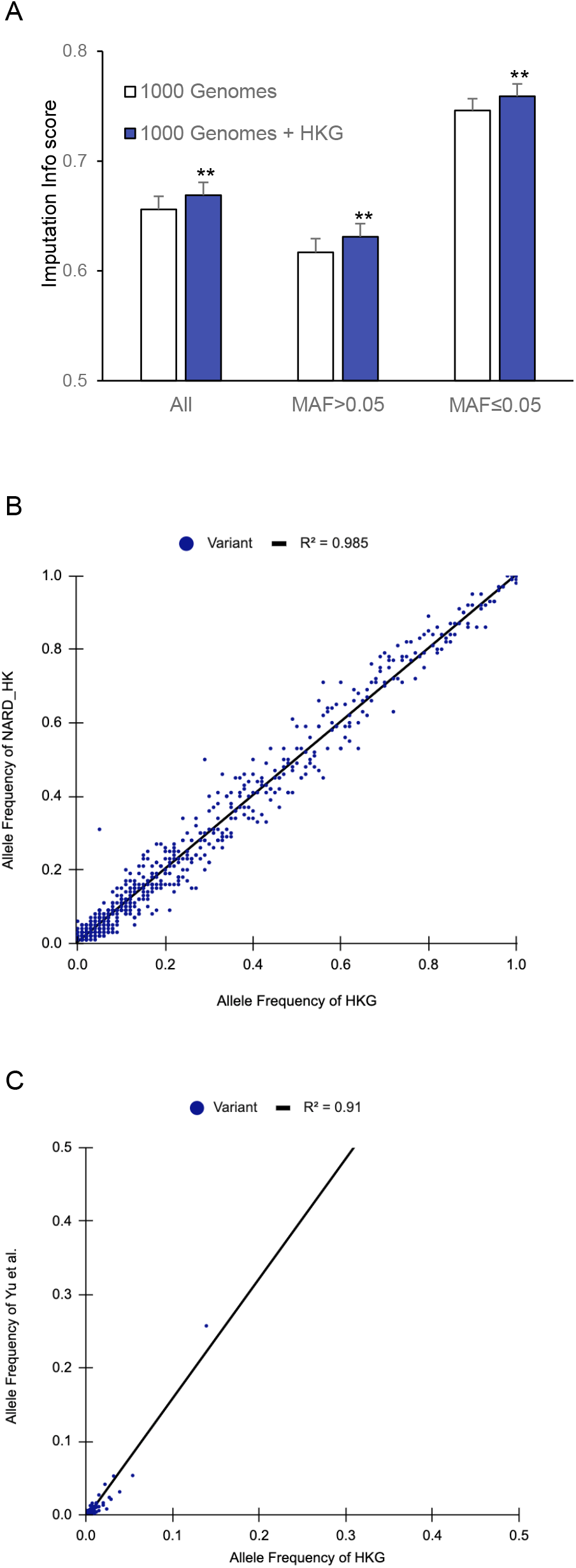
Validation of HKG variants by imputation and correlation analysis. (A) Imputation testing using the two reference panels: 1KGP and 1KGP + HKG. The average Info scores ± standard deviation error was based on 22 chromosomes. ** indicates the difference meets a significant level with p<0.01 of student’s T test. (B) Correlation analysis using AFs of variants in HKG and NARD_HK. (C) Correlation analysis using AFs of variants in HKG and Yu *et al*. 2021 reported actionable pharmacogenetic variants.

For correlation analysis, we obtained a limited number of Hong Kong samples from two existing studies and checked their variant consistency with HKG. The Northeast Asian Reference Database (NARD;(9)) involves 58 whole genome sequencing Hong Kong samples. It is the known database with the greatest number of variants (8,898,677 variants) from a Hong Kong population. The variants in these samples are hereafter abbreviated as NARD_HK. We found 17,224 variants in both NARD_HK and HKG. Figure 4B shows that allele frequencies in NARD_HK are well correlated (r^2^ = 0.985) with HKG, suggesting a strong positive linear association. Using Cook’s distance (> 0.0002), seven variants were identified to be common in NARD_HK but rare in HKG (Supplementary Table S9). Among them, only two variants, chr11:55639048C>T and chrX:71104307C>T, could be found in dbSNP, but none of them had a record of clinical significance. We also correlated the novel variants of HKG with the pharmacogenetic variants reported by Yu *et al*., 2021 (15). A linear relationship between their variant AFs was still observed (Figure 4C), but with a lower r^2^ of 0.909 due to the strong sampling bias towards pharmaceutical use in that study. These results highlight the effective correlation between HKG with other Hong Kong samples, hence they are a good representation of the local genetic diversity.

## Discussion

Among 612 search results of “Hong Kong Chinese variant” and “genotyping”/”WES”/”WGS” in PubMed (on Dec 14, 2020), 192 used samples collected in Hong Kong, but only 84 of them provided the count or frequency of the variants in individuals (Details can be found in Supplementary Table S10). As discussed above, there has long been a research gap in the development of a Hong Kong Cantonese-specific genomic database, which has hindered the regional and local development of genetic studies. Population-specific genomic data is undoubtedly one of the most valuable knowledge databases, which evolves with the ancestry and demography of a population (53). It allows us to trace the population origin or predict population susceptibility to future environmental changes (54). There is an increasing number of population-wide genetics studies (55,56), driven not only by the availability of resources, but also the related practical health care benefits. In this study, we illustrated the power of using WES as an alternative approach to describe the landscape of genetic variations to define the genomic characterizations of HKG. This holds the potential for novel gene characterizations at a lower cost than WGS (16). As HKG is the first public Hong Kong Cantonese variant database, it is the leading project to promote local genetic studies for research and clinical applications.

It is obvious that Hong Kong Cantonese consist mainly of Chinese Cantonese. The backbone of HKG is closest to Chinese CHB, CHS, and CDX populations sampled by 1KGP, with over 60% similarity. The majority of Hong Kong Cantonese share ancestry with CHS, and both have a similar effective population size. Our analysis also suggests sufficiently large uniqueness of HKG to justify the need to have a population-specific variant database. HKG is the first to demonstrate the potential association between ancestral composition and geographical distribution of Hong Kong Cantonese and provides evidence for the historical interpretation of Hong Kong Cantonese population migration, filling the gap of genetic diversity among Cantonese people. Interestingly, HKG also successfully captured a portion of non-Chinese diversity among the Hong Kong Cantonese. These samples might represent the South Asians who migrated to Hong Kong many years ago for historical reasons (1). This also suggests the comprehensiveness in our sampling.

Developments towards precision medicine are one of the most important medical challenges in recent years (57). Although a large proportion of variants identified in HKG are known variants, we updated the population-specific frequency and haplotype for 599 potentially pathogenic and druggable variants, based on the CIViC, PharmGKB and ClinVar annotations. The dynamics in population allele frequencies influences the functional interpretation of variants, which may assist local medical consultations and treatment decisions (57,58). HKG found 11,659 novel variants, 6.85% of which were high-impact variants. Since these novel variants had not been documented, biological associations were made only based on the corresponding gene and their association from disease databases. As different exomic variants could be mapped to the same coding genes, the biological associations should be carefully interpreted. Therefore, the HKG-specific disease-gene network presented in this study should only be regarded as predicted patterns and not to be taken as clinical advice. Instead, we encourage further investigation of possible genetically related health issues in the community, using HKG data as a steppingstone.

HKG is by far the largest public variant database for Hong Kong Cantonese. With its highly confident variant data, HKG can serve as a pioneering genomic resource region wide. As a part of southern Chinese, HKG also provides additional data for genetic studies of human traits in Chinese populations. The application effectiveness of our data is reflected in the imputation study and correlation analysis. Although HKG has already shown high discovery power to detect high-quality variants using high-depth exome data, the number of samples is still limited at this stage, and we believe that large-scale sampling in the future, such as HKGP, would be meaningful for further investigation.

## Data Availability

HKG is publicly available in the European Variation Archive (EVA) under study number PRJEB41688 (https://www.ebi.ac.uk/ena/browser/view/PRJEB41688). The genotype imputation can be freely performed at the HKG imputation server for the academic purpose at http://www.bio8.cs.hku.hk/HKGimputationServer.

## Ethics statement

This project was reviewed and approved by the University of Hong Kong Human Research Ethics Committee (HREC reference number: EA200067). This was a secondary data analysis project for sequencing protocol and bioinformatics algorithm development, and informed consent was not required by the Committee. All procedures were performed in accordance with relevant guidelines and regulations.

## Supplementary Materials

Supplementary Table S1-10.xlsx Supplementary Figure S1-5.pdf

## Reference

1. Carroll, J.M. (2007) A concise history of Hong Kong. Rowman & Littlefield Publishers.

2. Zhang, Z. and Ye, H. (2018) Mode of Migration, Age at Arrival, and Occupational Attainment of Immigrants from Mainland China to Hong Kong. Chinese Sociological Review, 50, 83–112.

3. Zhang, Z. and Wu, X. (2011) Social change, cohort quality and economic adaptation of Chinese immigrants in Hong Kong, 1991–2006. Asian and Pacific migration journal, 20, 1–29.

4. Siva, N. (2008). Nature Publishing Group.

5. Stephenson, J. (2008) 1000 Genomes Project. JAMA, 299, 755.

6. GenomeAsia, K.C. (2019) The GenomeAsia 100K Project enables genetic discoveries across Asia. Nature, 576, 106–111.

7. Siva, N. (2008) 1000 Genomes project. Nature Biotechnology, 26, 256–256.

8. Sherry, S.T. (2001) dbSNP: the NCBI database of genetic variation. Nucleic Acids Research, 29, 308–311.

9. Yoo, S.-K., Kim, C.-U., Kim, H.L., Kim, S., Shin, J.-Y., Kim, N., Yang, J.S.W., Lo, K.-W., Cho, B. and Matsuda, F. (2019) NARD: whole-genome reference panel of 1779 Northeast Asians improves imputation accuracy of rare and low-frequency variants. Genome medicine, 11, 1–10.

10. Wang, C.-C., Yeh, H.-Y., Popov, A.N., Zhang, H.-Q., Matsumura, H., Sirak, K., Cheronet, O., Kovalev, A., Rohland, N. and Kim, A.M. (2021) Genomic insights into the formation of human populations in East Asia. Nature, 591, 413–419.

11. Cao, Y., Li, L., Xu, M., Feng, Z., Sun, X., Lu, J., Xu, Y., Du, P., Wang, T. and Hu, R. (2020) The ChinaMAP analytics of deep whole genome sequences in 10,588 individuals. Cell research, 30, 717–731.

12. Zhang, P., Luo, H., Li, Y., Wang, Y., Wang, J., Zheng, Y., Niu, Y., Shi, Y., Zhou, H. and Song, T. (2020) NyuWa Genome Resource: Deep Whole Genome Sequencing Based Chinese Population Variation Profile and Reference Panel. bioRxiv.

13. Tiffin, P. and Ross-Ibarra, J. (2017) Advances and Limits of Using Population Genetics to Understand Local Adaptation: (Trends in Ecology & Evolution 29, 673-680; 2014). Trends Ecol. Evol., 32, 801–802.

14. Ashley, E.A. (2016) Towards precision medicine. Nat. Rev. Genet., 17, 507–522.

15. Yu, M.H.C., Chan, M.C.Y., Chung, C.C.Y., Li, A.W.T., Yip, C.Y.W., Mak, C.C.Y., Chau, J.F.T., Lee, M., Fung, J.L.F., Tsang, M.H.Y. et al.. (2021) Actionable pharmacogenetic variants in Hong Kong Chinese exome sequencing data and projected prescription impact in the Hong Kong population. PLOS Genetics, 17, e1009323.

16. Chou, J., Ohsumi, T.K. and Geha, R.S. (2012) Use of whole exome and genome sequencing in the identification of genetic causes of primary immunodeficiencies. Curr. Opin. Allergy Clin. Immunol., 12, 623–628.

17. Luo, R., Wong, Y.-L., Law, W.-C., Lee, L.-K., Cheung, J., Liu, C.-M. and Lam, T.-W. (2014) BALSA: integrated secondary analysis for whole-genome and whole-exome sequencing, accelerated by GPU. PeerJ, 2, e421.

18. Li, H., Handsaker, B., Wysoker, A., Fennell, T., Ruan, J., Homer, N., Marth, G., Abecasis, G., Durbin, R. and Genome Project Data Processing, S. (2009) The Sequence Alignment/Map format and SAMtools. Bioinformatics, 25, 2078–2079.

19. Institute, B. (2019) Picard toolkit. Broad Institute, GitHub repository.

20. McKenna, A., Hanna, M., Banks, E., Sivachenko, A., Cibulskis, K., Kernytsky, A., Garimella, K., Altshuler, D., Gabriel, S. and Daly, M. (2010) The Genome Analysis Toolkit: a MapReduce framework for analyzing next-generation DNA sequencing data. Genome research, 20, 1297–1303.

21. Consortium, t.I.H. and †The International HapMap, C. (2003) The International HapMap Project. Nature, 426, 789–796.

22. Mills, R.E., Pittard, W.S., Mullaney, J.M., Farooq, U., Creasy, T.H., Mahurkar, A.A., Kemeza, D.M., Strassler, D.S., Ponting, C.P., Webber, C. et al.. (2011) Natural genetic variation caused by small insertions and deletions in the human genome. Genome Res., 21, 830–839.

23. Lek, M., Karczewski, K.J., Minikel, E.V., Samocha, K.E., Banks, E., Fennell, T., O’Donnell-Luria, A.H., Ware, J.S., Hill, A.J., Cummings, B.B. et al.. (2016) Analysis of protein-coding genetic variation in 60,706 humans. Nature, 536, 285–291.

24. Karczewski, K.J., Francioli, L.C., Tiao, G., Cummings, B.B., Alföldi, J., Wang, Q., Collins, R.L., Laricchia, K.M., Ganna, A., Birnbaum, D.P. et al.. (2020) The mutational constraint spectrum quantified from variation in 141,456 humans. Nature, 581, 434–443.

25. Bergström, A., McCarthy, S.A., Hui, R., Almarri, M.A., Ayub, Q., Danecek, P., Chen, Y., Felkel, S., Hallast, P., Kamm, J. et al.. (2020) Insights into human genetic variation and population history from 929 diverse genomes. Science, 367.

26. Zheng, X., Levine, D., Shen, J., Gogarten, S.M., Laurie, C. and Weir, B.S. (2012) A high-performance computing toolset for relatedness and principal component analysis of SNP data. Bioinformatics, 28, 3326–3328.

27. Zheng, X., Gogarten, S.M., Lawrence, M., Stilp, A., Conomos, M.P., Weir, B.S., Laurie, C. and Levine, D. (2017) SeqArray-a storage-efficient high-performance data format for WGS variant calls. Bioinformatics, 33, 2251–2257.

28. Browning, S.R. and Browning, B.L. (2007) Rapid and accurate haplotype phasing and missing-data inference for whole-genome association studies by use of localized haplotype clustering. Am. J. Hum. Genet., 81, 1084–1097.

29. Browning, B.L. and Browning, S.R. (2016) Genotype Imputation with Millions of Reference Samples. Am. J. Hum. Genet., 98, 116–126.

30. Nakatsuka, N., Moorjani, P., Rai, N., Sarkar, B., Tandon, A., Patterson, N., Bhavani, G.S., Girisha, K.M., Mustak, M.S., Srinivasan, S. et al.. (2017) The promise of discovering population-specific disease-associated genes in South Asia. Nat. Genet., 49, 1403–1407.

31. Purcell, S., Neale, B., Todd-Brown, K., Thomas, L., Ferreira, M.A.R., Bender, D., Maller, J., Sklar, P., de Bakker, P.I.W., Daly, M.J. et al.. (2007) PLINK: A Tool Set for Whole-Genome Association and Population-Based Linkage Analyses. The American Journal of Human Genetics, 81, 559–575.

32. Alexander, D.H. and Lange, K. (2011) Enhancements to the ADMIXTURE algorithm for individual ancestry estimation. BMC Bioinformatics, 12, 246.

33. Francis, R.M. (2017) pophelper: an R package and web app to analyse and visualize population structure. Molecular Ecology Resources, 17, 27–32.

34. Cvijović, I., Good, B.H. and Desai, M.M. (2018) The Effect of Strong Purifying Selection on Genetic Diversity. Genetics, 209, 1235–1278.

35. Jeroncic, A., Memari, Y., Ritchie, G.R.S., Hendricks, A.E., Kolb-Kokocinski, A., Matchan, A., Vitart, V., Hayward, C., Kolcic, I., Glodzik, D. et al.. (2016) Whole-exome sequencing in an isolated population from the Dalmatian island of Vis. European Journal of Human Genetics, 24, 1479–1487.

36. Wang, Q., Pierce-Hoffman, E., Cummings, B.B., Alföldi, J., Francioli, L.C., Gauthier, L.D., Hill, A.J., O’Donnell-Luria, A.H., Genome Aggregation Database Production, T., Genome Aggregation Database, C. et al. (2020) Landscape of multi-nucleotide variants in 125,748 human exomes and 15,708 genomes. Nat. Commun., 11, 2539.

37. Online Mendelian Inheritance in Man, O. (2021) McKusick-Nathans Institute of Genetic Medicine, Johns Hopkins University (Baltimore, MD).

38. Janeway, C., Travers, P., Walport, M. and Schlomchik, M. (2001) Immunobiology 5 : the Immune System in Health and Disease. Garland Science.

39. Nait Saada, J., Kalantzis, G., Shyr, D., Cooper, F., Robinson, M., Gusev, A. and Palamara, P.F. (2020) Identity-by-descent detection across 487,409 British samples reveals fine scale population structure and ultra-rare variant associations. Nat. Commun., 11, 6130.

40. Naseri, A., Tang, K., Geng, X., Shi, J., Zhang, J., Shakya, P., Liu, X., Zhang, S. and Zhi, D. (2021) Personalized genealogical history of UK individuals inferred from biobank-scale IBD segments. BMC Biol., 19, 1–12.

41. Snapshot of Hong Kong Population.

42. Li, Q., Ma, R. and Zhang, M. (2018) XRCC1 rs1799782 (C194T) polymorphism correlated with tumor metastasis and molecular subtypes in breast cancer. OncoTargets and Therapy, 11, 8435–8444.

43. Correia, C.T., Almeida, J.P., Santos, P.E., Sequeira, A.F., Marques, C.E., Miguel, T.S., Abreu, R.L., Oliveira, G.G. and Vicente, A.M. (2010) Pharmacogenetics of risperidone therapy in autism: association analysis of eight candidate genes with drug efficacy and adverse drug reactions. Pharmacogenomics J., 10, 418–430.

44. Zhang, J.-P., Lencz, T., Zhang, R.X., Nitta, M., Maayan, L., John, M., Robinson, D.G., Fleischhacker, W.W., Kahn, R.S., Ophoff, R.A. et al.. (2016) Pharmacogenetic Associations of Antipsychotic Drug-Related Weight Gain: A Systematic Review and Meta-analysis. Schizophr. Bull., 42, 1418–1437.

45. Houston, J.P., Kohler, J., Bishop, J.R., Ellingrod, V.L., Ostbye, K.M., Zhao, F., Conley, R.R., Hoffmann, V.P. and Fijal, B.A. (2012) Pharmacogenomic Associations With Weight Gain in Olanzapine Treatment of Patients Without Schizophrenia. J. Clin. Psychiatry, 73, 1077–1086.

46. Fang, P., He, J.-Y., Han, A.-X., Lan, T., Dai, D.-P., Cai, J.-P. and Hu, G.-X. (2017) Effects of CYP2C19 Variants on Fluoxetine Metabolism in vitro. Pharmacology, 100, 91–97.

47. Xu, R.-A., Gu, E.-M., Liu, T.-H., Ou-yang, Q.-G., Hu, G.-X. and Cai, J.-P. (2018) The effects of cytochrome P450 2C19 polymorphism on the metabolism of voriconazole in vitro. Individ. Differ. Res., 11, 2129–2135.

48. Wang, H., Kim, R.A., Sun, D., Gao, Y., Wang, H., Zhu, J. and Chen, C. (2011) Evaluation of the effects of 18 non-synonymous single-nucleotide polymorphisms of CYP450 2C19 on in vitro drug inhibition potential by a fluorescence-based high-throughput assay. Xenobiotica, 41, 826–835.

49. Jahan, R., Macha, M.A., Rachagani, S., Das, S., Smith, L.M., Kaur, S. and Batra, S.K. (2018) Axed MUC4 (MUC4/X) aggravates pancreatic malignant phenotype by activating integrin-β1/FAK/ERK pathway. Biochim. Biophys. Acta Mol. Basis Dis., 1864, 2538–2549.

50. Du, X. and Gomez, C.M. (2018) Spinocerebellar [corrected] Ataxia Type 6: Molecular Mechanisms and Calcium Channel Genetics. Adv. Exp. Med. Biol., 1049, 147–173.

51. Government announces appointments to Hong Kong Genome Institute.

52. GeneCards Human Gene, D. ZAN Gene - GeneCards.

53. Nielsen, R., Akey, J.M., Jakobsson, M., Pritchard, J.K., Tishkoff, S. and Willerslev, E. (2017) Tracing the peopling of the world through genomics. Nature, 541, 302.

54. Zeberg, H. and Pääbo, S. (2020) The major genetic risk factor for severe COVID-19 is inherited from Neanderthals. Nature, 587, 610–612.

55. (2015) The UK10K project identifies rare variants in health and disease. Nature, 526, 82–90.

56. Al-Ali, M., Osman, W., Tay, G.K. and AlSafar, H.S. (2018) A 1000 Arab genome project to study the Emirati population. J. Hum. Genet., 63, 533–536.

57. Olivier, M., Asmis, R., Hawkins, G.A., Howard, T.D. and Cox, L.A. (2019) The Need for Multi-Omics Biomarker Signatures in Precision Medicine. Int. J. Mol. Sci., 20.

58. Carrasco-Ramiro, F., Peiró-Pastor, R. and Aguado, B. (2017) Human genomics projects and precision medicine. Gene Ther., 24.

