## Supplementary Figures for "HKG: An open genetic variant database of 205 Hong Kong Cantonese exomes"

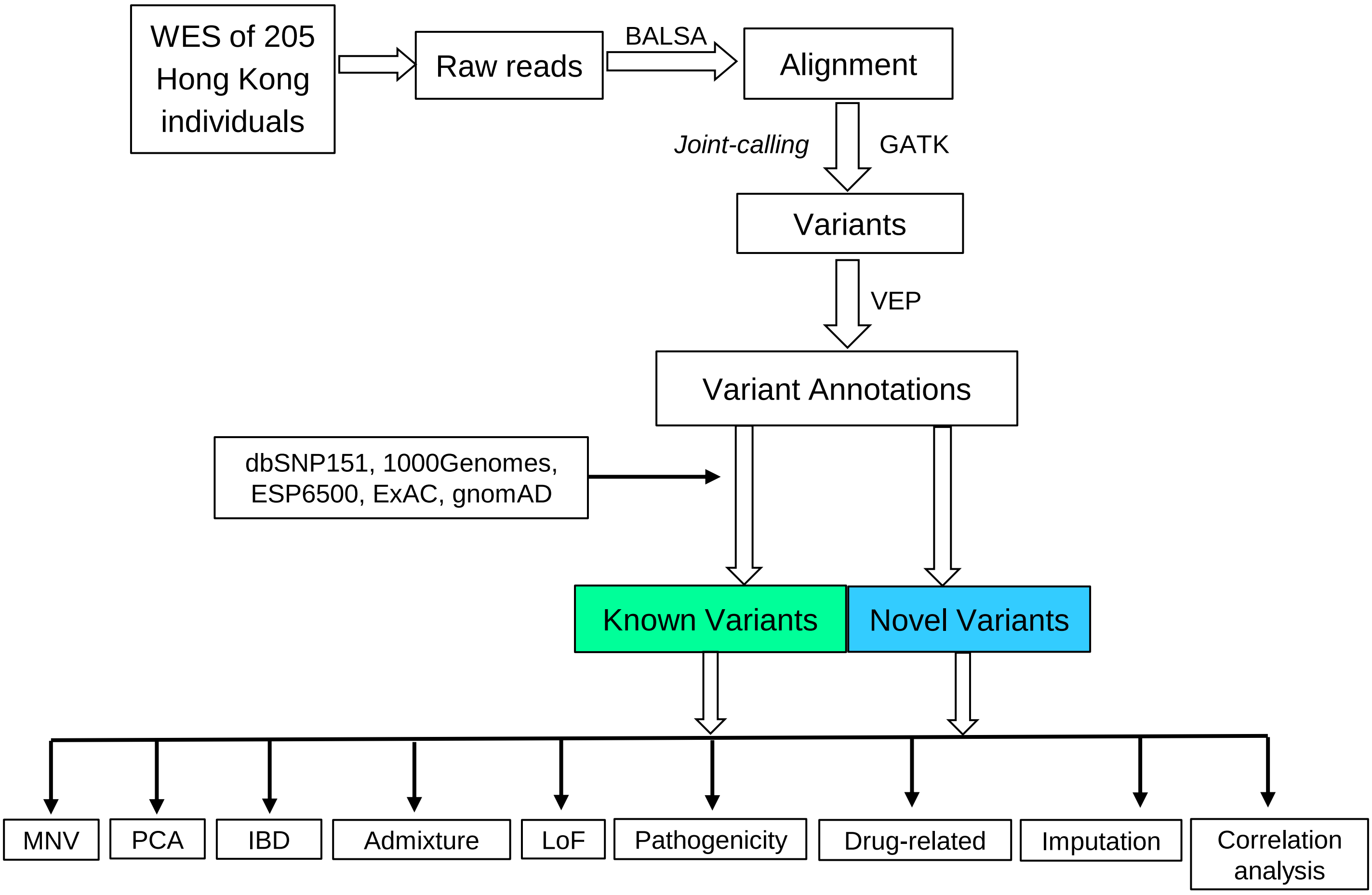


**Fig. S1.** An overview of discovery and characterization of genetic variants using the WES data of 205 Hong Kong individuals. After alignment of raw reads, we adopted joint-calling, quality control and filtering methods to call variants according to the Genome Analysis Toolkit (GATK) procedure. Using Variant Effect Predictor (VEP) of Ensembl, the identified variants were annotated into different consequence subsets. Five main public databases were used to divide the variants obtained from HKG into known and novel components for downstream characterization of HKG.

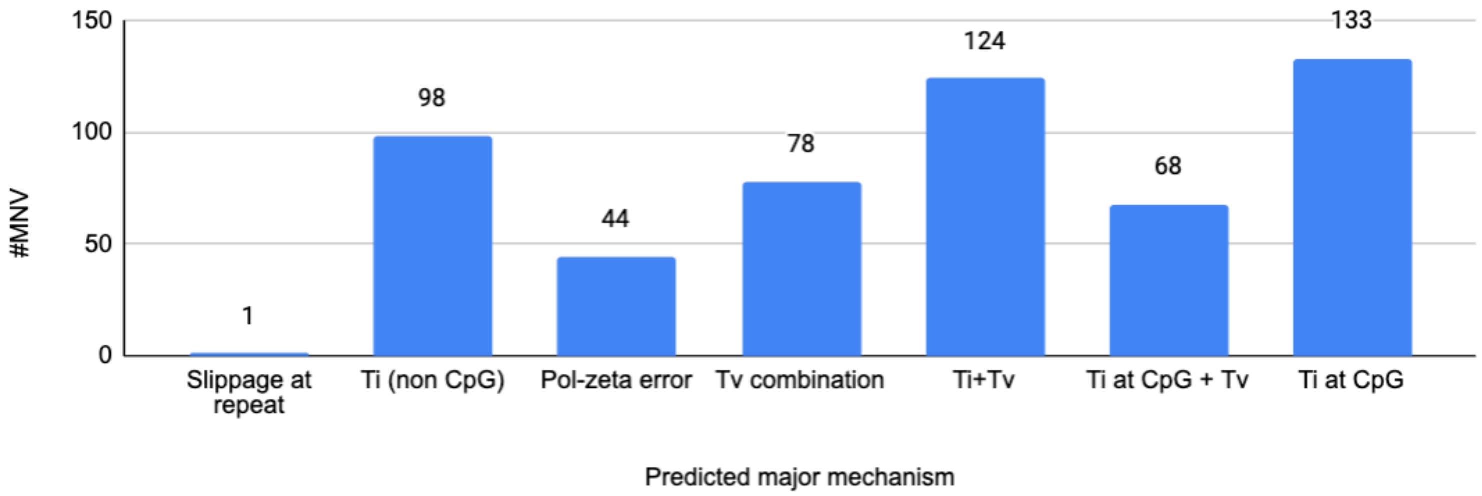


**Fig. S2.** Distribution of MNV with different patterns. Slippage at repeat: polymerase slippage at repeat junctions; Ti (non CpG): transition at non CpG site; Pol-zeta error: replication error introduced by DNA polymerase zeta; Tv combination: two transversions; Ti + Tv: one transition at non-CpG site and one transversion; Ti at CpG + Tv: one transition at CpG site and one transversion; and Ti at CpG: two transitions at CpG site.

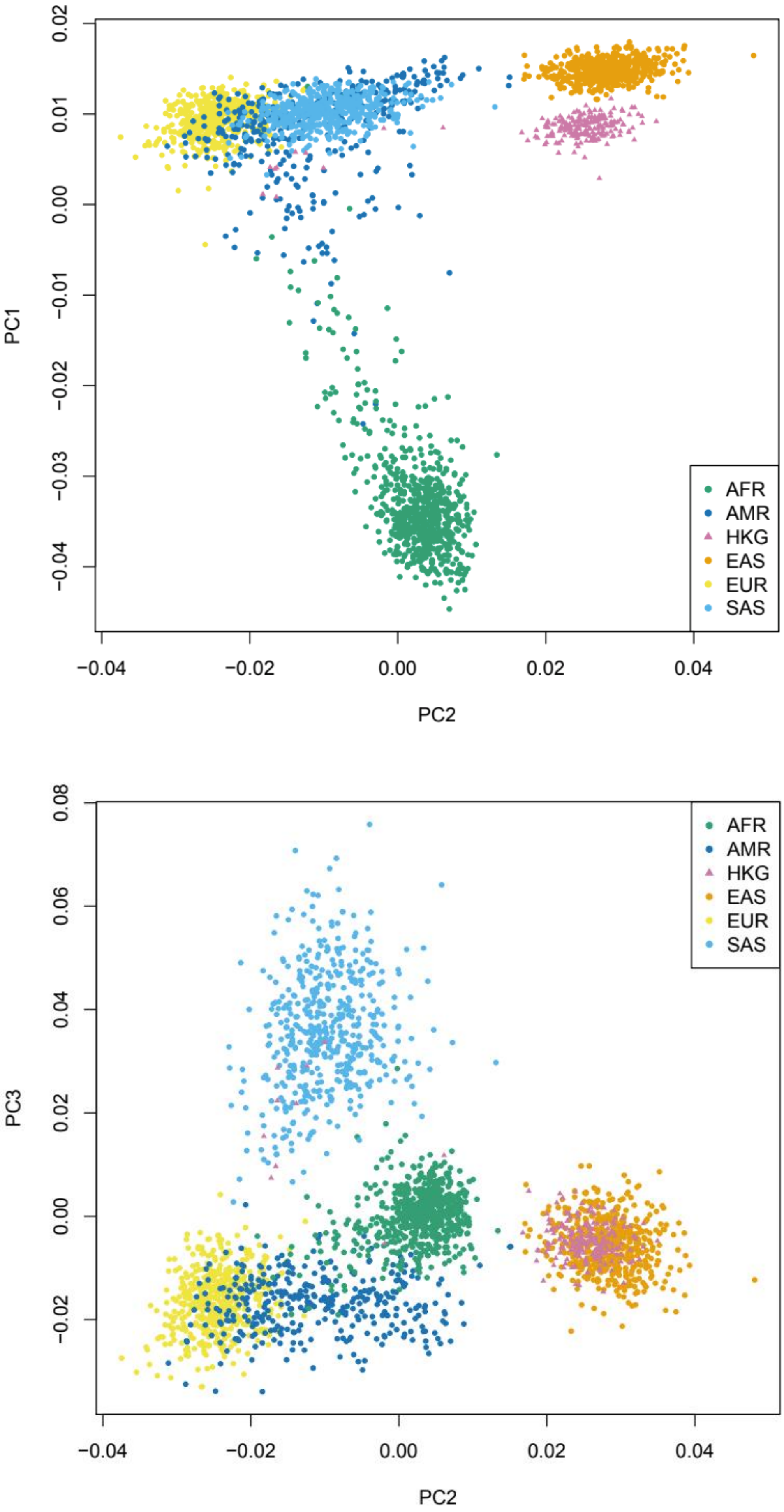


**Fig. S3.** PCA of HKG and five populations of 1000 Genomes AFR, AMR, EAS, EUR and SAS. AFR: Africans; AMR: Admixed Americans; EAS: East Asians; EUR: Europeans; and SAS: South Asians.

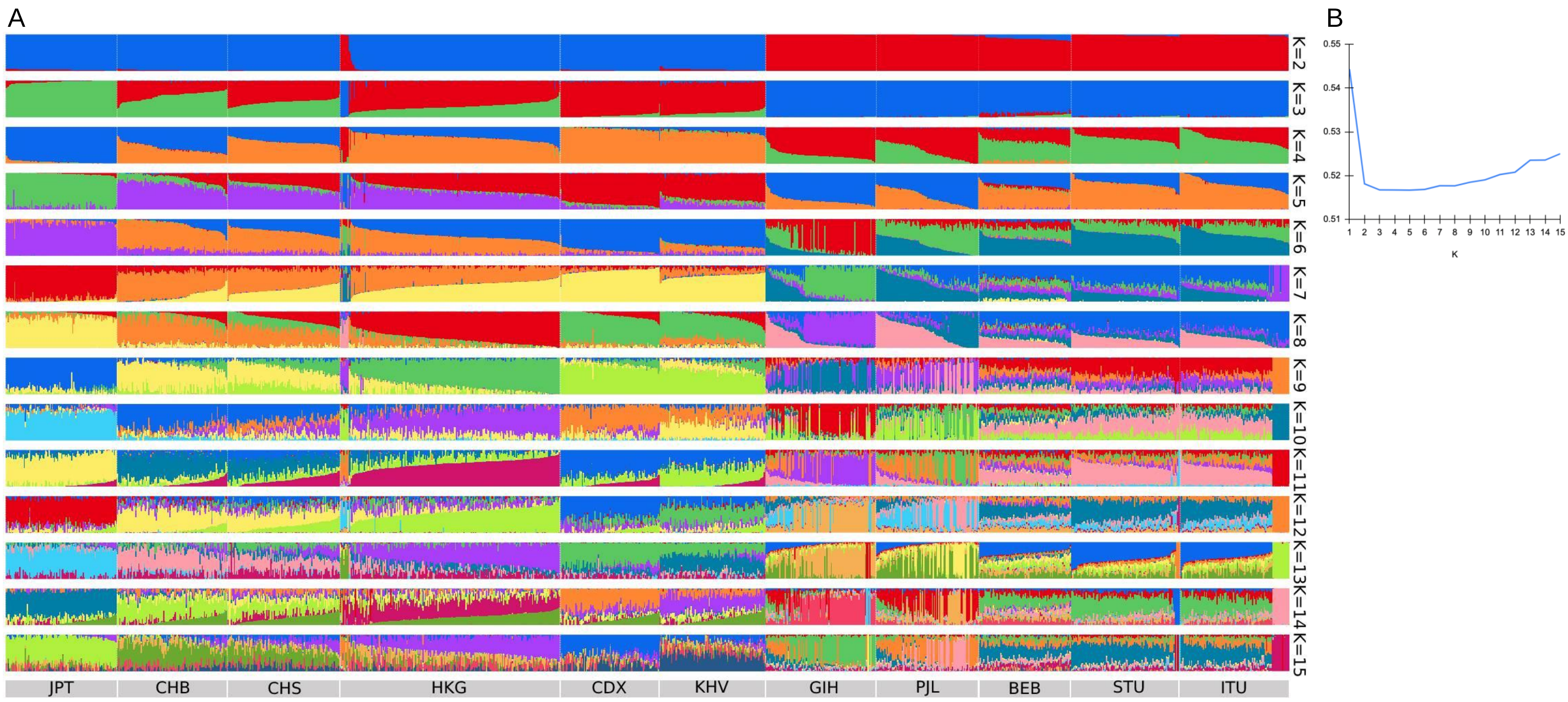


**Fig. S4.** Ancestral population composition of HKG compared with other East Asian and South Asian populations. A) the ADMIXTURE analysis of HKG samples with five East Asia and six South Asia populations obtained from 1KGP performed with K = 2 to K = 15. JPT: Japanese in Tokyo, Japan; CHB: Han Chinese in Beijing, China; CHS: Han Chinese South, China; CDX: Chinese Dai in Xishuangbanna, China; KHV: Kinh in Ho Chi Minh City, Vietnam; GIH: Gujarati Indians in Houston, Texas, United States; PJL: Punjabi in Lahore, Pakistan; BEB: Bengali in Bangladesh; STU: Sri Lankan Tamil in the UK; and ITU: Indian Telugu in the U.K.. B) The cross-validation (CV) of the ADMIXTURE analysis showed K = 5 best explained the dataset.

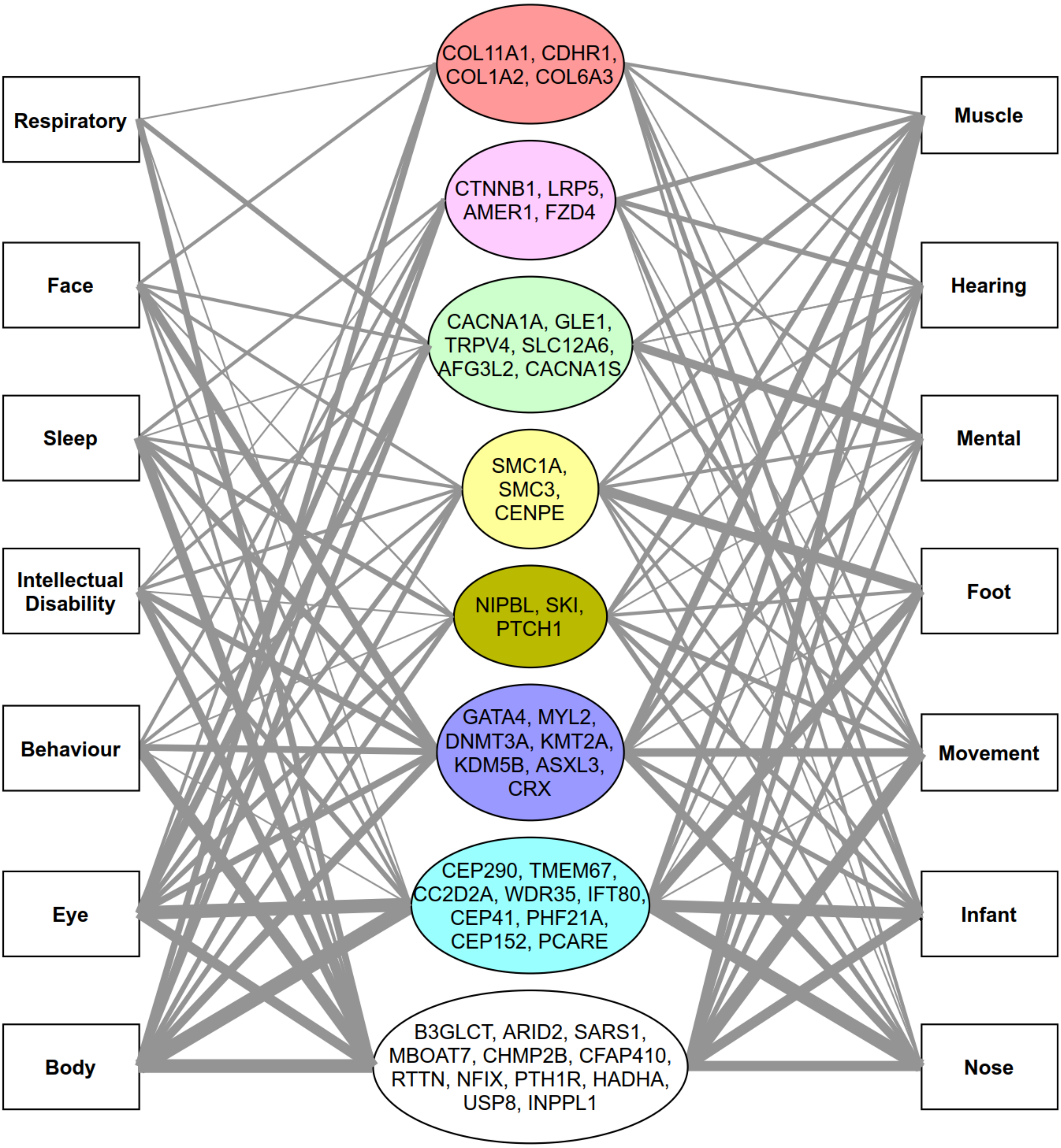


**Fig. S5.** The disease-gene network of novel high impact variants in HKG. Constructed based on 634 genes of novel high impact variants in HKG which were not found in other Chinese populations. The elliptical nodes represent these disease-associated genes with enrichment (i.e., p-value < 0.01; FDR < 0.25) affecting different biological processes including: orange (cell adhesion), brown (development), yellow (cell cycle), green (transmembrane transport), pink (Wnt signaling), light blue (cilium assembly) and purple (transcriptional or epigenetic modification). Rectangular nodes represent different disease types based on DisGeNET gene sets. The width of edges indicates the number of genes that involve this association.
